# Pathogenicity and selective constraint on variation near splice sites

**DOI:** 10.1101/256636

**Authors:** Jenny Lord, Giuseppe Gallone, Patrick J. Short, Jeremy F. McRae, Holly Ironfield, Elizabeth H. Wynn, Sebastian S. Gerety, Liu He, Bronwyn Kerr, Diana S. Johnson, Emma McCann, Esther Kinning, Frances Flinter, I. Karen Temple, Jill Clayton-Smith, Meriel McEntagart, Sally Ann Lynch, Shelagh Joss, Sofia Douzgou, Tabib Dabir, Virginia Clowes, Vivienne P. M. McConnell, Wayne Lam, Caroline F. Wright, David R. FitzPatrick, Helen V. Firth, Jeffrey C. Barrett, Matthew E. Hurles, on behalf of the Deciphering Developmental Disorders study

## Abstract

Mutations which perturb normal pre-mRNA splicing are significant contributors to human disease. We used exome sequencing data from 7,833 probands with developmental disorders (DD) and their unaffected parents, as well as >60,000 aggregated exomes from the Exome Aggregation Consortium, to investigate selection around the splice site, and quantify the contribution of splicing mutations to DDs. Patterns of purifying selection, a deficit of variants in highly constrained genes in healthy subjects and excess *de novo* mutations in patients highlighted particular positions within and around the consensus splice site of greater functional relevance. Using mutational burden analyses in this large cohort of proband-parent trios, we could estimate in an unbiased manner the relative contributions of mutations at canonical dinucleotides (73%) and flanking non-canonical positions (27%), and calculated the positive predictive value of pathogenicity for different classes of mutations. We identified 18 patients with likely diagnostic *de novo* mutations in dominant DD-associated genes at non-canonical positions in splice sites. We estimate 35-40% of pathogenic variants in non-canonical splice site positions are missing from public databases.

## Introduction

Pre-mRNA splicing in humans is mediated by the major and minor spliceosomes, highly dynamic, metalloenzyme complexes comprised of five key small nuclear RNAs (snRNA), along with over 100 protein components and accessory molecules(Brody and Abelson 1985; Hang et al. 2015; Scotti and Swanson 2016). Accurate recruitment and function of the spliceosome is reliant on a plethora of cis-acting regulatory elements encoded within the pre-mRNA itself. Whilst our understanding of the underlying mechanistic processes regulating splicing has greatly increased in recent years, our ability to predict whether or not a mutation will affect splicing remains limited. However, with estimates that up to 50% of monogenic disease-causing variants may affect splicing(Teraoka et al. 1999; Ars et al. 2000), a better understanding and more coherent approach to interpretation of variants affecting splicing is badly needed(Cartegni et al. 2002; Baralle and Buratti 2017). With a plethora of *in silico* splicing pathogenicity predictors available, there is little consensus on what a “gold standard” for splicing pathogenicity prediction would be(Houdayer et al. 2012; Jian et al. 2014b; Tang et al. 2016). Whilst many of these methods perform well within the canonical splice site dinucleotides (CSS, the two highly-conserved bases flanking the acceptor and donor sites), their utility for other splice relevant regions is less clear(Tang et al. 2016). In the clinical setting, often multiple algorithms and expert judgment are used to predict pathogenicity, while for large scale research projects, classification of variants is often binary, with CSS mutations typically classified as likely splice affecting, whilst mutations in other splicing regulatory components are typically overlooked(Iossifov et al. 2014; Wright et al. 2015; Deciphering Developmental Disorders 2017). Previous attempts to estimate the relative contribution of pathogenic variants at non-canonical splice sites were based on collating diverse published datasets of pathogenic variants (Lewandowska 2013) or data submitted to databases of clinically interpreted variation (Krawczak et al. 2007) and are therefore sensitive to the inherent heterogeneity and biases of such data, especially given the inevitable subjectivity involved in clinical interpretation of this class of variation. Both clinical and research interpretation of potential splice-disrupting variants lacks a robust quantitative foundation.

Utilising large scale exome sequencing data from 13,750 unaffected parents recruited as part of the Deciphering Developmental Disorders (DDD) project(Wright et al. 2015) and >60,000 aggregated exomes from the Exome Aggregation Consortium (ExAC) (Lek et al. 2016), we explore selective constraint around splice regions across a set of 148,244 stringently defined exons well covered (median coverage >15X at both CSS) across the DDD cohort (see Methods). Since selection is driven by a number of factors, including monogenic developmental disorders (DD), as a complementary, disease-based approach, we analyse enrichment of *de novo* mutations (DNMs) in DDD probands in the same regions. We provide an unbiased, exome wide view of the signatures of selection and the relative contribution of pathogenic splice altering mutations between the CSS and other, near splice positions.

## Results

### Signatures of purifying selection around the splice site

Since purifying selection acts to keep deleterious alleles rare, population variation data can be used to identify and assess the relative strengths of signals of purifying selection. To assess selective constraint acting on positions around the canonical splice site we used the Mutability Adjusted Proportion of Singletons (MAPS) metric(Lek et al. 2016) (a measure for inferring the degree of selection robust to local variance in mutation rate) in 13,750 unaffected parents enrolled in the DDD study as well as >60,000 aggregated exomes from ExAC (Figure 1a). The canonical splice acceptor and donor dinucleotides show a clear signal of purifying selection in both datasets.

**Figure 1.**
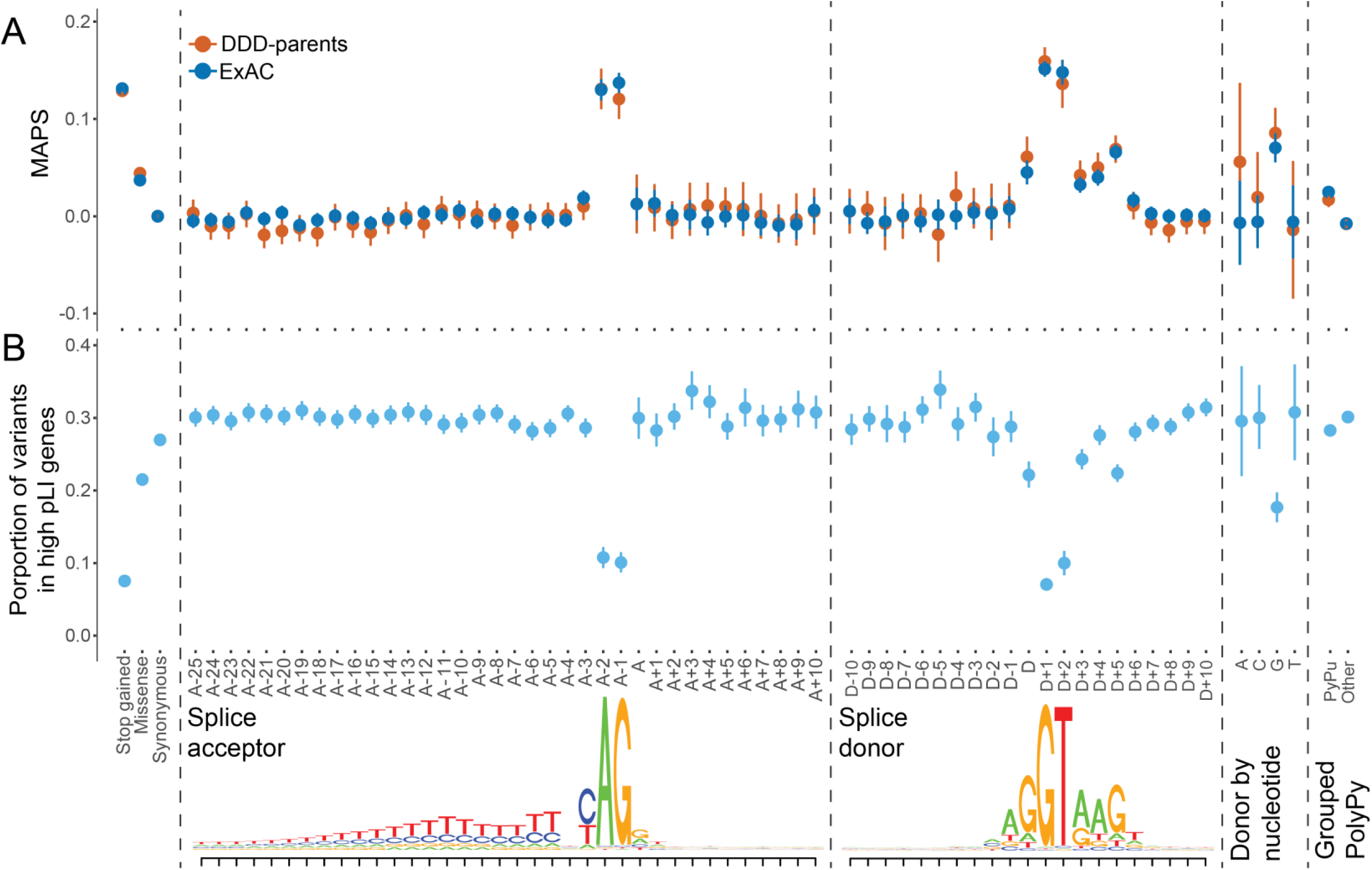
Signals of purifying selection around splice sites. A. Selective constraint across splicing region in 13,750 unaffected parents of DDD probands and >60,000 aggregated exomes from ExAC. Mutability adjusted proportion of singletons (MAPS) with 95% confidence intervals (CI) shown for Ensembl’s Variant Effect Predictor (VEP) annotated exonic sites, extended splice acceptor and splice donor regions, the last base of the exon, split by reference nucleotide, and grouped sites in the polypyrimidine tract region (PolyPy), split by changes from a pyrimidine to a purine (PyPu) vs all other changes. B. Proportion of variants with 95% CI in 13,750 unaffected parents of DDD probands which fall within genes with high probability of loss of function intolerance (pLI >0.9) across VEP annotated exonic sites, extended splice acceptor and splice donor regions, the last base of the exon, split by reference nucleotide, and grouped sites in the polypyrimidine tract region, split by changes from a pyrimidine to a purine (PyPu) vs all other changes. Lower panel shows splice acceptor and splice donor consensus sequences, based on our exons of interest.

Outside of the CSS, other positions clearly show a signal of purifying selection beyond the background level, including the donor site (last base of the exon, which is particularly pronounced when the reference allele is G (Figure 1a)), and the intronic positions proximal to the canonical donor site, peaking at the don+5 position, which exhibits a signal of purifying selection intermediate between missense and stop-gained variants. Although no sites within the polypyrimidine tract (PolyPy) show a signal of purifying selection individually, when these sites are grouped together (Methods) and stratified by changes from a pyrimidine to a purine (PyPu), versus all other changes, there is a clear difference between the two types of variants, with PyPu changes exhibiting an increased signal of purifying selection when compared to non-PyPu changes (bootstrap p < 0.001, Figure 1a, Supplemental Fig S1).

### Deficit of splicing variants in highly constrained genes in healthy individuals

We also examined the distribution of variants of different classes among genes that are known to be under different levels of selective constraint. Highly constrained genes should contain fewer deleterious variants than less constrained genes. We investigated the proportion of variants observed in the 13,750 unaffected parents which fell within highly constrained genes (probability of loss of function (LoF) intolerance (pLI, (Lek et al. 2016)) > 0.9) in our splicing regions of interest (Figure 1b). In the near splice positions at which the highest MAPS values were seen (CSS, donor, donor+5), we also observed a stronger depletion of variants in high pLI genes within the unaffected parents, again supporting the potential pathogenicity of variants at these positions. The proportion of parental variants in high pLI genes also recapitulates the signals of purifying selection seen in the MAPS analyses with regard to the donor position split by reference allele (Figure 1b) and the PolyPy region (Figure 1b, Supplemental Fig. S1), with the lowest proportions in high pLI genes observed for sites with the highest MAPS values.

### Assessing the significance of mutational burden for different classes of splicing mutations

We identified 871 autosomal high confidence DNMs (non-synonymous consequences excluded) within canonical and near-splice regions of interest well covered by exome data in the 7,833 probands, allowing us to test for enrichment of DNMs relative to expectations based on a trinucleotide null model of mutation rate (Samocha et al. 2014) across different sets of genes (DD-associated with dominant or recessive mechanisms, and non-DD associated, see Methods). Across recessive DD and non-DD associated genes, no enrichment of DNMs beyond the null expectation was observed (Figure 2a). In dominant DD genes, a significant cumulative excess of DNMs was noted across the full splicing region (Poisson p (false discovery rate (FDR) adjusted) = 1.33×10^-14^, fold enrichment = 3.47), which remained significant upon exclusion of the canonical dinucleotide positions (Poisson p (FDR adjusted) = 0.0035, fold enrichment = 1.86). Individually, the four canonical splice site positions each showed a significant (10-27 fold) excess of DNMs (Poisson p (FDR adjusted), fold enrichment: acc-2 = 4.22×10^-12^, 26.6; acc-1 = 3.43×10^-8^, 16.6; don+1 = 1.33×10^-14^, 20.1; don+2 = 0.004, 10.0), as did the don+5 site (9.7×10^-5^, 9.29). The similar level of enrichment between don+5 and don+2 implies these positions harbour comparable proportions of splice disrupting mutations. No individual positions within the PolyPy region showed an individual excess of DNMs, however, when the positions were considered cumulatively and split between PyPu and non-PyPu changes (Figure 2b), an excess of DNMs was observed in the PyPu group for dominant DD genes (fold enrichment = 3.46), although this was not significant at an FDR of 5% (Poisson p (FDR corrected) = 0.086).

**Figure 2.**
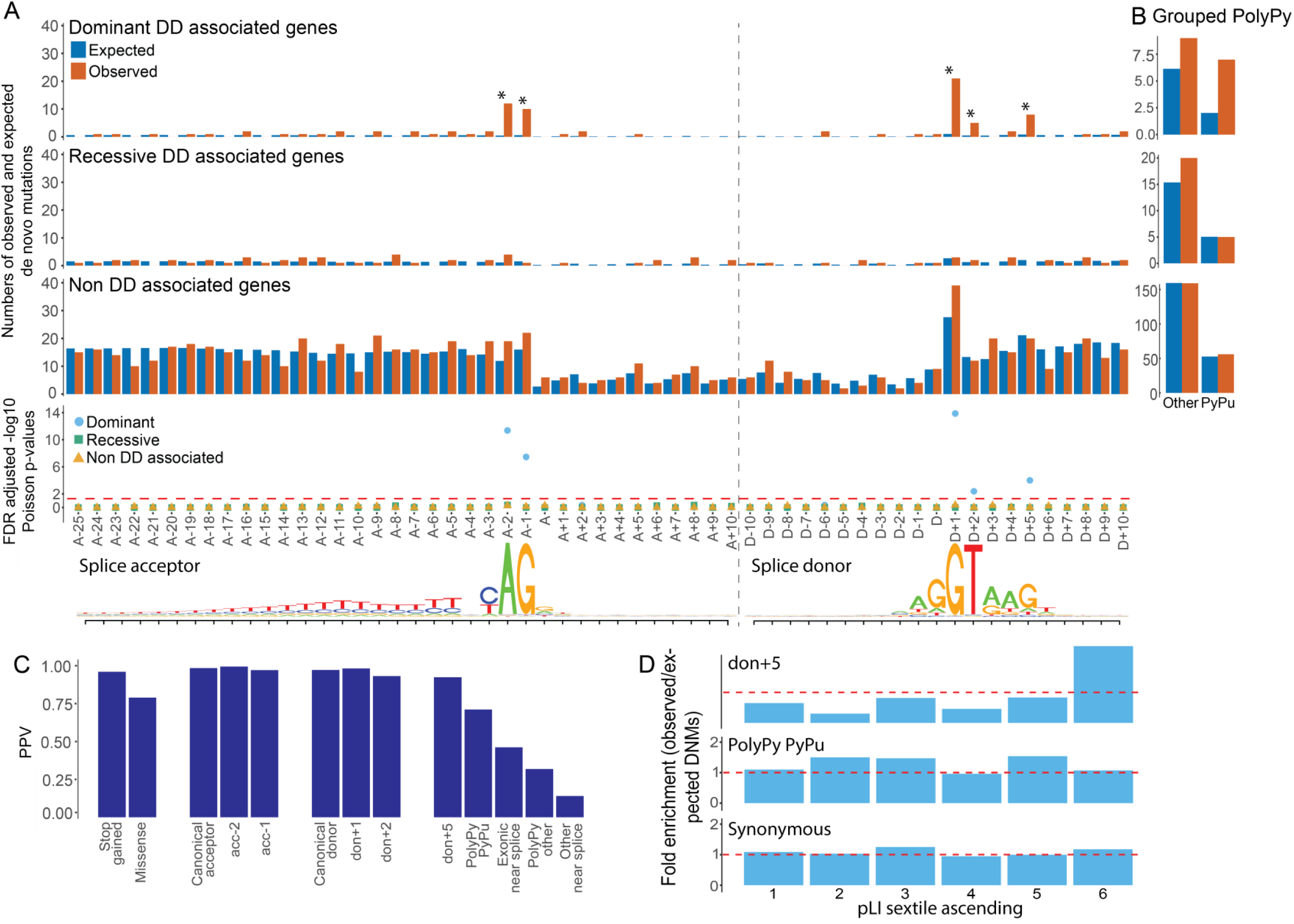
*De novo* mutations around splice sites. Enrichment of *de novo* mutations (DNMs) across the splicing region in 7,833 DDD probands A. Numbers of observed and expected DNMs across splicing region, in known dominant and recessive DD genes, as well as non-DD associated genes, with FDR corrected Poisson p-values. Splice acceptor and splice donor consensus sequences shown below, as in Figure 1. B. Aggregation of observed and expected numbers of DNMs in the polypyrimidine tract (PolyPy) region, with changes from a pyrimidine to a purine (“PyPu”) and all other changes shown separately for known dominant and recessive DD genes, as well as non-DD associated genes. C. Positive predictive values (PPVs) for *de novo* mutations in dominant DD-associated genes in positions across the splicing region, as well as VEP annotated stop gained and missense changes, calculated from observed and expected numbers of DNMs. D. Enrichment (observed/expected) of *de* novo mutations (DNMs) by gene probability of loss of function intolerance (pLI), split in to sextiles for donor+5, pyrimidine to purine PolyPy, and synonymous sites. pLI scores encompassed by each sextile: 1 = 5.36E-91 - 0.000000605, 2 = 0.000000609 - 0.000558185, 3 = 0.000559475 - 0.027905143, 4 = 0.027908298 - 0.377456159, 5 = 0.377491926 - 0.919495985, 6 = 0.91955878 - 1.

When the same analysis was performed for dominant genes in subsets of the DDD cohort with (n=1417) and without (n=3364) robust diagnoses from the standard diagnostic protocol, which only assesses splicing mutations at the CSS (Supplemental Fig. S2), the enrichment within the diagnosed subset was confined to the CSS (Poisson p (FDR adjusted), fold enrichment: CSS = 1.33−10^-14^, 69.74; other positions = 0.658, 1.82), whilst in the undiagnosed subset, the opposite pattern was observed (Poisson p (FDR adjusted), fold enrichment: CSS = 1, 0; other positions = 0.012, 2.21), with the don+5 site showing the greatest enrichment (16.18, Poisson p (FDR adjusted) = 5.35×10^-5^).

These results are highly concordant with the signatures of purifying selection identified using the MAPS metric, and the deficit of parental variants in high pLI genes, providing multiple independent lines of evidence that mutations in positions outside of the CSS can disrupt normal splicing.

### Estimating positive predictive values for different classes of splice mutation

We used the fold-enrichment of the numbers of observed DNMs in dominant DD genes in the DDD cohort over the number expected under the null mutation model to calculate positive predictive values (PPVs) for groupings of near splice site positions. We compared these with PPVs for other, more commonly disease-associated variant classes within the same exons of the same genes (Figure 2c). We observe minor differences in PPV for the individual positions of the canonical acceptor and donor sites, with don+2 showing the lowest PPV at 0.90, which is approximately the same as for the don+5 position (PPV 0.89). Variants within the PolyPy region which change a pyrimidine for a purine have a PPV of 0.71, which is below the PPV for missense mutations (0.79), but still predicts a substantive number of pathogenic mutations arising from disruption of the PolyPy.

Despite the modest number of observed DNMs used to make these PPV estimates, we see striking concordance with the population based metrics described above (MAPS and deficit of splicing variants in high pLI genes in unaffected parents of DDD patients – Supplemental Fig. S3), suggesting these estimates are robust.

We looked at the distribution of observed DNMs in genes with respect to their probability of being LoF intolerant (using the pLI metric(Lek et al. 2016), Figure 2d). For synonymous variants, we observed no significant enrichment of DNMs in high pLI genes. For don+5 mutations, there is a clear excess of DNMs in genes most likely to be intolerant to LoF mutations in the DDD cohort, further supporting the likely pathogenicity of mutations in these positions. For the PolyPy PyPu mutations, although there is a nominally significant enrichment of DNMs in general, this does not show a significant skew towards high pLI genes in our cohort.

### Identifying diagnostic non-canonical splice mutations

After exclusion of probands with likely diagnostic protein-coding or canonical splice site variants, 38 DNMs in our near splice site positions of interest in dominant DD genes were identified. The clinical phenotypes of patients carrying these mutations were reviewed by a consultant clinical geneticist, blinded to the precise mutation and PPVs estimated above, and the patient’s recruiting clinician, to assess the phenotypic similarity between the proband and the disorder expected from a LoF mutation in that gene. The 38 variants were classified as likely diagnostic (Table 1), or unlikely diagnostic/unknown (Supplemental Table S1), depending on the strength of phenotypic similarity. Phenotypic information for probands with likely diagnostic variants is given in Supplemental Table S2, and pathogenicity prediction scores for the SNVs in Supplemental Table S3. The clinical review resulted in 18 variants (47%) being classified as likely diagnostic, highly concordant with the number predicted from the overall PPV of non-canonical sites of 46%, moreover, a higher proportion of likely diagnostic variants were classified at sites with higher PPVs (Pearson’s correlation coefficient = 0.91, p = 0.033) (Figure 3). With 48 CSS DNMs observed within the same exons in our probands, we estimate that 73% (95% CI: 60-82%) of disease causing splice disrupting DNMs occur within the CSS, while 27% (95% CI: 18-39%) are in non-canonical, near-splice positions.

**Figure 3.**
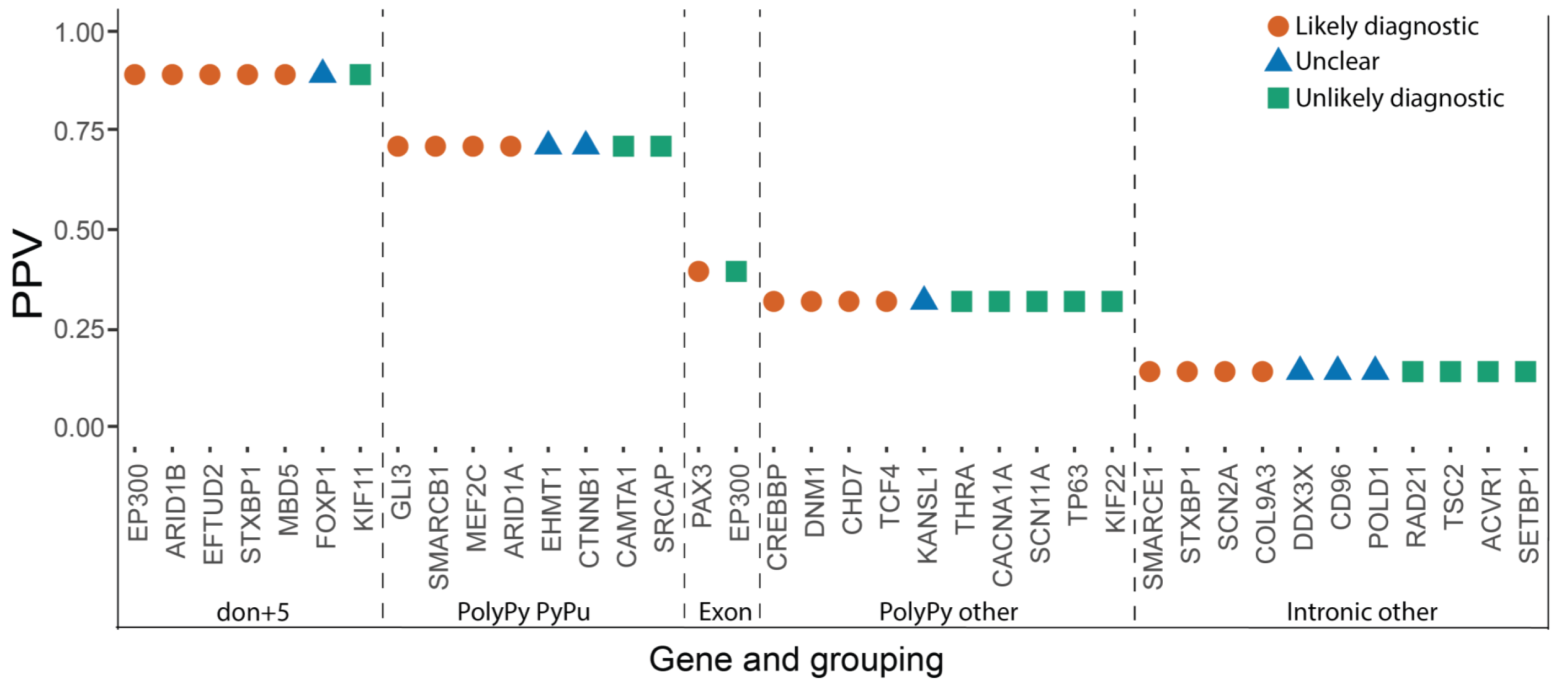
Clinical classifications of non-canonical near splice *de novo* mutations. Relationship between clinical classifications of 38 splice region *de novo* mutations (DNMs) in undiagnosed DDD probands and positive predictive values (PPVs) calculated using observed and expected numbers of DNMs in 7,833 probands.

Eight DNMs were selected for functional validation via a minigene vector system, including six likely diagnostic PolyPy variants, a PolyPy variant of uncertain clinical significance, and a likely diagnostic don+5 variant, where both the phenotype of the patient and that associated with the gene (*MBD5*) are nonspecific, along with two negative controls (untransmitted variants identified in unaffected parents within the same PolyPys as test variants). For six of the variants selected for validation, differences in splicing between the reference and mutant constructs were observed (Supplemental Fig. S4a and S4b). One of the likely diagnostic PolyPy mutations, the PolyPy mutation of uncertain significance, and both negative controls showed no difference in splicing between the reference and mutant constructs (Supplemental Fig. S4c and S4d).

**Table 1.**
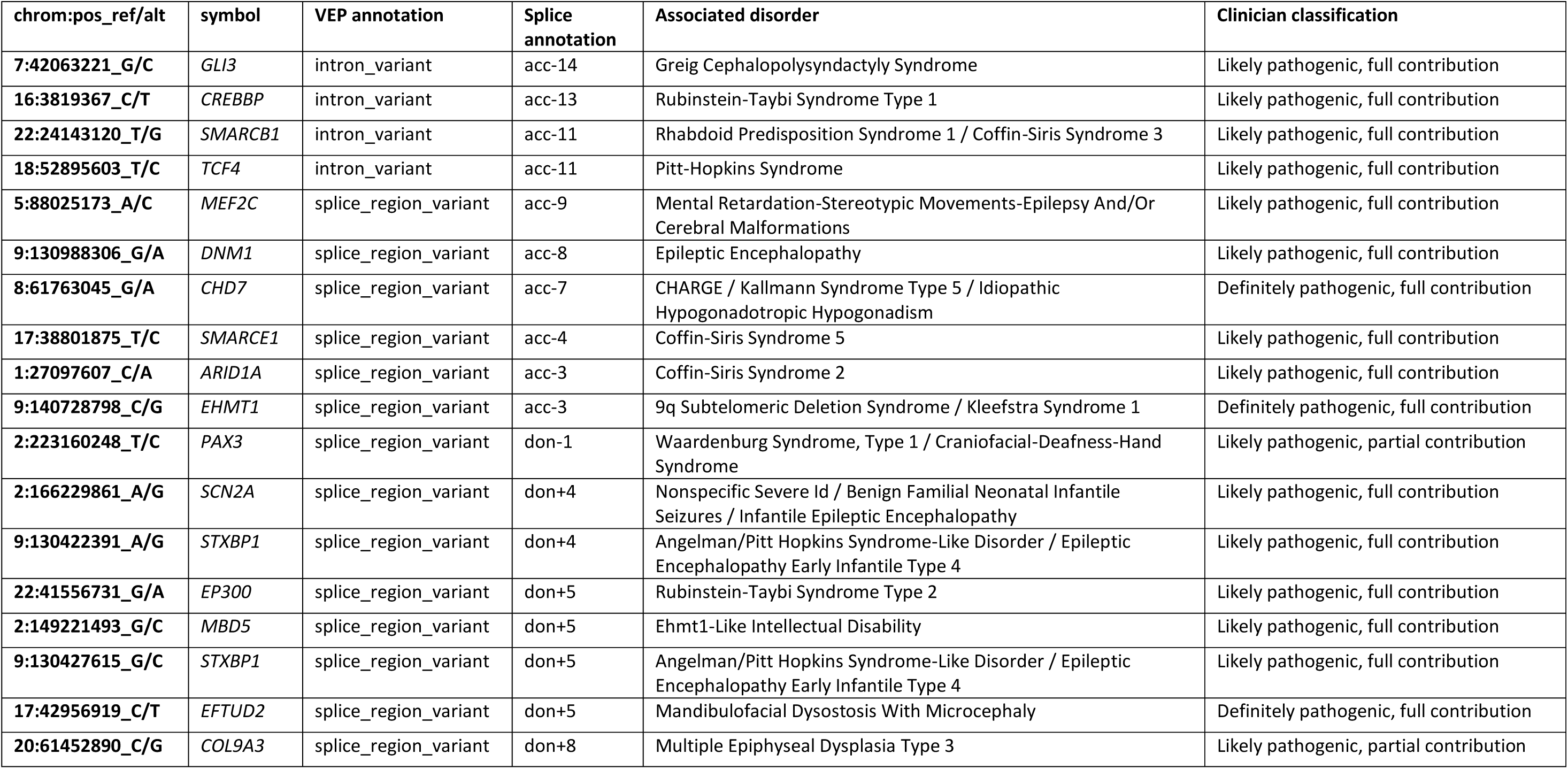
Diagnostic *de novo* mutations in non-canonical dinucleotide near splice positions. Variant and proband information for 18 *de novo* likely diagnostic splice region variants identified in previously undiagnosed DDD probands in known dominant DD-associated genes (hg19 coordinates).

### Assessing splicing pathogenicity prediction tools

The population genetic metrics of purifying selection and mutation enrichment metric for pathogenicity that we have derived provide an orthogonal approach to assessing the accuracy of splicing pathogenicity prediction tools, compared to the standard approach of assessing classification accuracy for clinically interpreted variants. We assessed four splicing pathogenicity prediction tools: two recently published genome-wide ensemble learning methods: AdaBoost and RandomForest, Spidex (based on deep learning trained on RNA sequencing data), and the longer standing, widely used MaxEntScan (MES) (Yeo and Burge 2004; Jian et al. 2014a; Xiong et al. 2015).

We divided the scores from each prediction tool, plus CADD(Kircher et al. 2014), into 20 equal-sized bins to facilitate cross-method comparability. We calculated the MAPS for each bin of each of the scoring metrics for the splicing variants observed in the 13,750 DDD unaffected parents, and saw a strong positive correlation between pathogenicity metric and MAPS for all tools (Figure 4). AdaBoost had the highest absolute MAPS value for the top scoring bin, suggesting that it is best able to identify variants under the strongest purifying selection. The proportion of variants in the unaffected parents falling in genes with pLI > 0.9 broadly recapitulates this pattern, with fewer variants in high pLI genes in the highest scoring brackets for all metrics (Supplemental Fig. S5). We then looked at the distribution of scores for each tool for the 83 splicing DNMs observed in DDD probands in autosomal dominant DD-associated genes which were covered by all five scoring systems to compare performance of the metrics on mutations more likely to have a deleterious impact on splicing, with the expectation that these potentially damaging variants would be scored highly by the metrics, giving high values of area under the curve (AUC, Figure 5). Again, all metrics performed well, with the majority of DNMs being classified in the most deleterious score brackets. Here AdaBoost gave the highest AUC value, suggesting it weighted these likely damaging variants as more deleterious than the other metrics comparatively. Interestingly, when CSS positions were removed from the analysis, AdaBoost remained the tool with the highest AUC. The largest reduction in the AUC metric was seen for Spidex and CADD, indicating these tools may be least informative for positions outside of the CSS. Upon removal of the CSS positions from the analyses of MAPS and deficit of parental variants in high pLI genes, similar results were revealed, with the highest AdaBoost scores retaining strong signals of purifying selection but a marked reduction in signal from the highest Spidex scores (Supplemental Fig. S6 and Supplemental Fig. S7).

**Figure 4.**
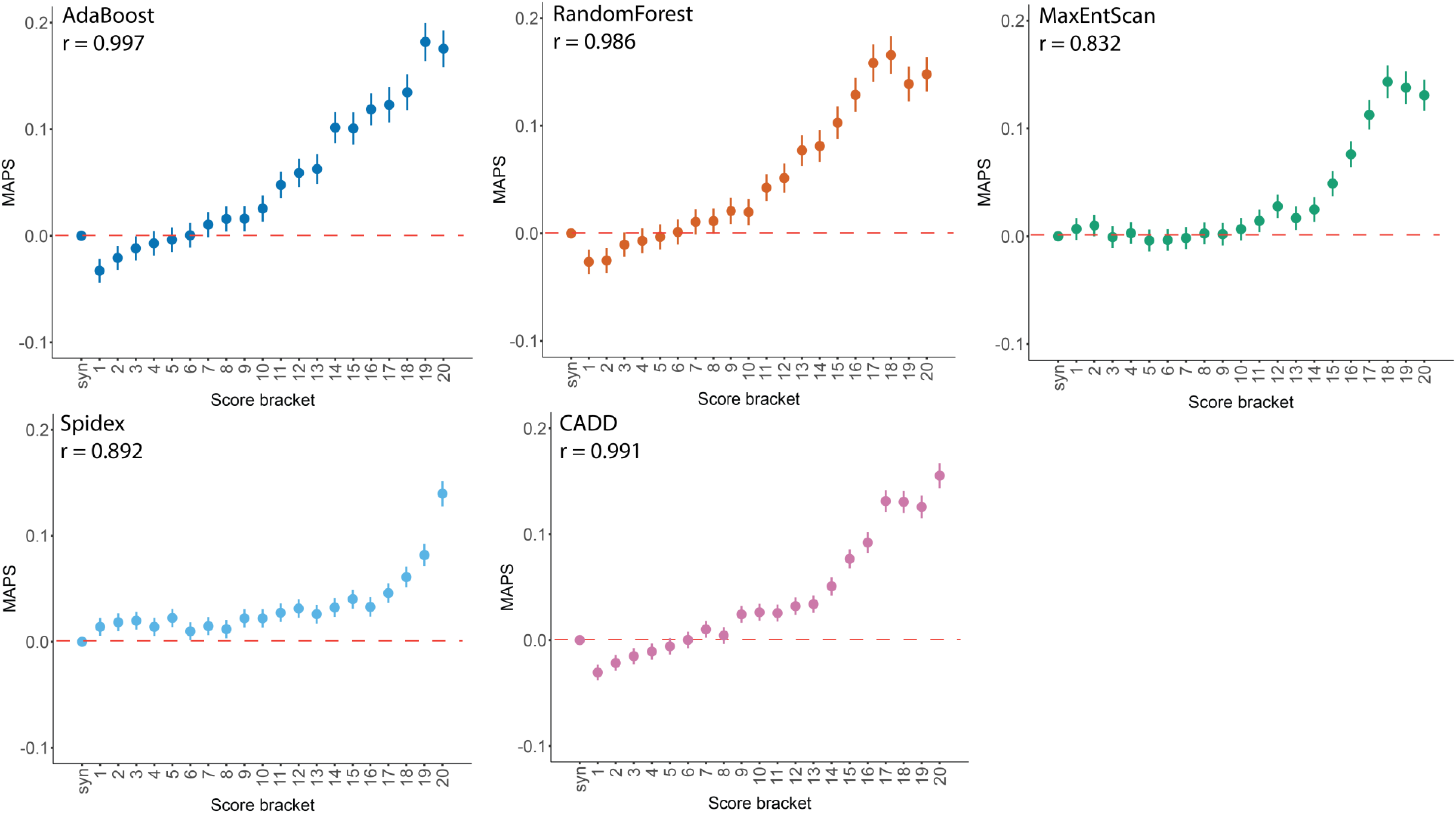
Selective constraint and pathogenicity scores. Mutability adjusted proportion of singletons (MAPS), with 95% CI, calculated for pathogenicity score brackets (least to most severe) in 13,750 unaffected parents from the DDD project, with Spearman correlation coefficient.

**Figure 5.**
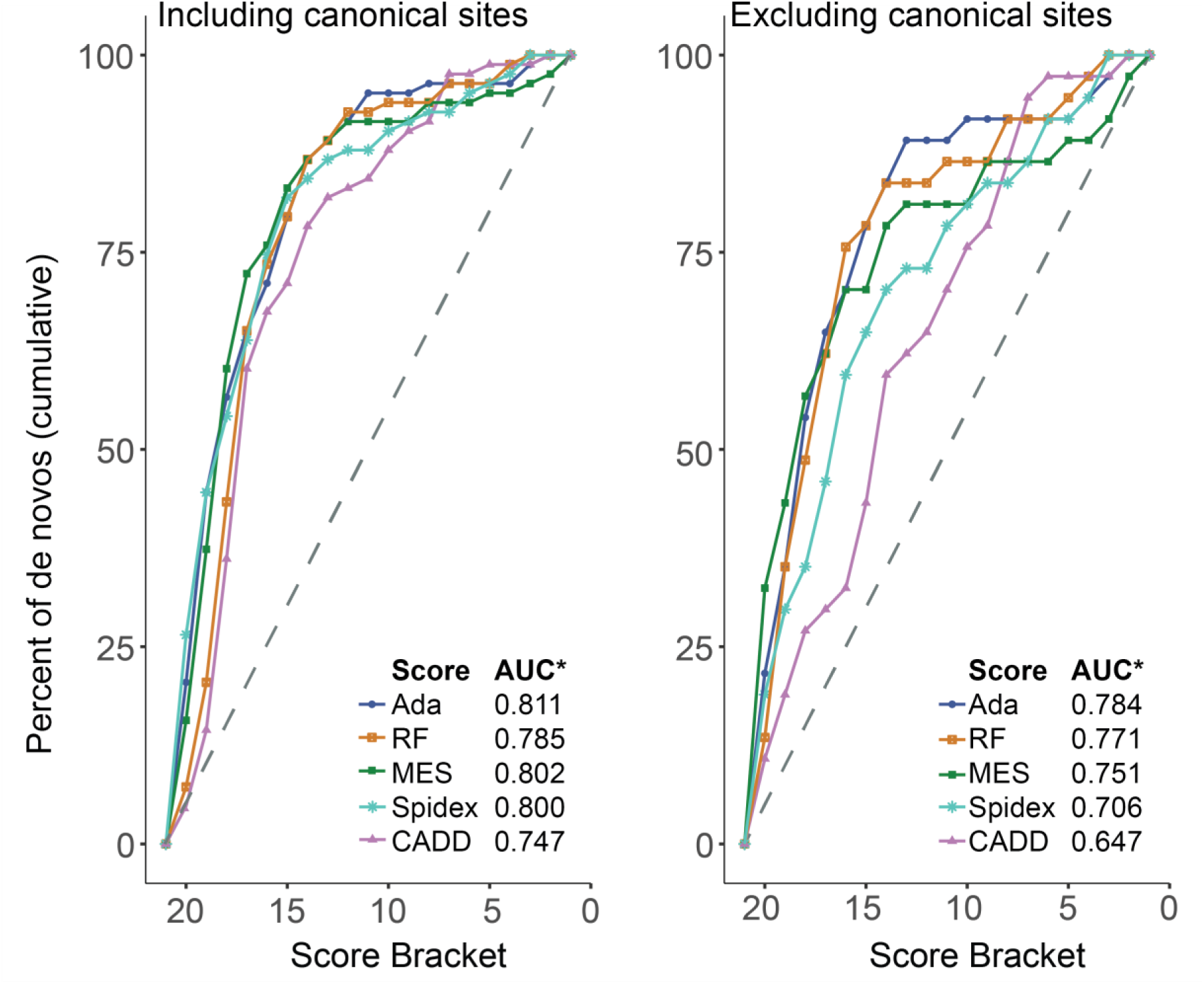
Pathogenicity scores for observed near splice site *de novo* mutations. Cumulative percentage of *de novo* mutations (DNMs) in known dominant DD genes with decreasing pathogenicity score bracket, shown with canonical splice site positions included (left) and excluded (right). * AUC = area under curve

Taken together, these data show a strong relationship between the considered splicing pathogenicity scoring systems and the general landscape of purifying selection on splicing control, but demonstrate that the utility of these systems in identifying likely diagnostic variants is limited outside of the CSS.

## Discussion

Our study represents a large, unbiased exploration of the perturbation of splicing by genetic variation in near-splice regions, with complementary signals of selection observed through two different population-based analyses. Selection can be driven by many factors, including monogenic disease resulting in reduced reproductive fitness. DDs represent the largest single class of monogenic diseases. With a clear enrichment of DNMs demonstrated by previous studies of DDs (Deciphering Developmental Disorders 2017), analysis of such DNMs provided an independent, disease-based approach, complementary to our population-based analyses. Concordance in results from the different approaches indicates the robustness of our conclusions.

Our analyses, taken together, suggest the pathogenic contribution of non-canonical splice positions has been under-appreciated. We estimate that around 27% (95% CI: 18-39%) of splice disrupting pathogenic mutations within the DDD cohort are in non-canonical positions. In sites with pathogenic or likely pathogenic clinical significance in ClinVar(Landrum et al. 2016) overlapping with our splicing positions of interest (and with non-synonymous consequences removed), we found 83.5% of variants fell within canonical positions, with just 16.5% in non-canonical positions. When adjusted for number of submissions as a proxy for allele count, this figure was 17.5%, perhaps indicative that recurrence strengthens evidence of pathogenicity. Both of these values are significantly below our estimate of 27% (Fisher’s Exact p = 1.22×10^-15^ and p < 2.2×10^-16^ respectively), suggesting under-ascertainment of non-canonical splicing variants by around 35-40% in clinical databases, despite a growing understanding of the importance of such sites in splicing regulation(Kircher et al. 2014; Ferreira et al. 2016; Soukarieh et al. 2016; Cummings et al. 2017; Ito et al. 2017; Soemedi et al. 2017; Ke et al. 2018).

Estimates of the relative contribution of canonical and non-canonical splice site mutations are sparse in the literature. These estimates are also typically based on clinically interpreted variants and so are likely to be biased by the accuracy of current clinical practices. When comparing canonical and non-canonical mutations within the Human Gene Mutation Database (HGMD, based on variation described in publications, Krawczak *et al*.(Krawczak et al. 2007) stated canonical mutations accounted for 64% of mutations at donor sites and 77.4% of mutations at acceptor sites, giving an estimated non-canonical contribution of ˜30% overall (consistent with our data), while data taken from Caminsky *et al*.(Caminsky et al. 2014) put this estimate at around 43% (above our upper bound). These values are closer to our 27% estimate than to the ClinVar proportion of ˜17%, despite our approach focussing on DNMs and dominant disorders, whereas the other two studies did not discriminate on mode of inheritance and included recessive disorders, which can also be caused by non-canonical splicing mutations(Basel-Vanagaite et al. 2013; Brunham et al. 2015) and exonic variants. Our findings highlight the complementarity of assessing the clinical importance of non-canonical splice variants both through the traditional approach based on clinically-interpreted variation accrued through diagnostic practice and through unbiased approaches that leverage population variation and unbiased models of germline mutation.

Our analysis of non-canonical splice position mutations did not include exonic missense variants(Teraoka et al. 1999; Ars et al. 2000; Krawczak et al. 2007), nor did it explicitly include branchpoints(Maslen et al. 1997; Di Leo et al. 2004; Crotti et al. 2009; Aten et al. 2013), splicing enhancers and suppressors(Lorson et al. 1999; Liu et al. 2001) or deep intronic mutations(Cummings et al. 2017; Vaz-Drago et al. 2017). Detecting splice disrupting variants at these sites is even more challenging, as despite recent efforts(Corvelo et al. 2010; Wang and Wang 2014; Mercer et al. 2015; Badr et al. 2016; Taggart et al. 2017), comprehensive catalogues of all branchpoints and exonic and intronic splicing enhancers and silencers are currently unavailable, algorithms that predict the impact of mutations at such sites are not highly accurate, and some of these sites are not covered by exome sequencing (the greater utilisation of whole genome sequencing will allow greater opportunity to find and assess the contributions of more distal splice disrupting variants). As such, our estimate of the contribution of non-canonical splicing position mutations is likely to be a lower bound. Thus our estimate of 35-40% under-ascertainment in clinical databases may be conservative, and the true extent of missed diagnoses may be even higher.

The size of the available datasets determines our power to detect signals of selection and enrichment of DNMs, so although we could demonstrate signal at splice-important non-canonical positions, there may be other positions with more subtle signals of selection which we lacked the power to detect with current sizes of datasets. Collation of yet larger datasets will help to identify these sites, using the same methodology applied here.

The nature of many developmental disorders makes obtaining RNA samples from relevant tissues of patients (i.e. neural tissue) acutely problematic, so we investigated the effects on splicing of several of the potentially diagnostic DNMs using a minigene vector system. We were able to demonstrate changes to splicing for five out of six likely diagnostic PolyPy variants as well as the likely diagnostic don+5 variant, supporting the clinical interpretation based on clinical phenotype. We did not observe an effect on splicing for one likely diagnostic PolyPy variant, and one PolyPy variant of uncertain clinical significance. Although the accuracy of minigene assays when compared with patient RNA is generally high(Bonnet et al. 2008; Thery et al. 2011; van der Klift et al. 2015), known limitations of the system (e.g. lack of full endogenous genetic context(Baralle et al. 2006; Sangermano et al. 2017), and sensitivity to cell type utilised(Lastella et al. 2006)) mean we cannot definitively state that the effects seen in the minigene assay would be the same in the full genetic, developmental and cellular context within the patient.

We envisage that greater appreciation of the importance of near splice site mutations will increase diagnostic yields, as well as providing increased power for the detection of new genetic associations, both within the field of rare disease and beyond. We highlight two challenges to improving detection of pathogenic non-canonical splice site mutations.

First, many commonly used *in silico* tools for annotating the likely functional impact of variants do not discriminate between different non-CSS positions with very different likelihoods of being pathogenic. Moreover, commonly used annotation tools differ in the ways in which variants are annotated, with splicing variants displaying the highest level of disagreement between tools(McCarthy et al. 2014). This highlights the need for a more consistent and evidence based annotation of splicing variants. Of the positions shown in our analyses to be most damaging, don+5 sites are annotated by VEP(McLaren et al. 2016) and SnpEff(Cingolani et al. 2012) as “splice_region_variant”, while most positions of the PolyPy are annotated as intronic, so are potentially easily overlooked. With Annovar’s(Wang et al. 2010) default settings, only the CSSs are flagged as splicing variants, although with both Annovar and SnpEff, the user can optionally extend the region to be annotated as splice variants. We note that Ensembl have recently implemented a VEP plugin which allows greater granularity in splice region annotation (https://github.com/Ensembl/VEP_plugins/blob/release/88/SpliceRegion.pm), including annotating the don+5 and other near-donor positions, as well as the PolyPy region. This type of increased granularity of splicing annotation should facilitate consideration of these variants in future studies.

Second, current tools that predict the pathogenicity of non-CSS mutations have limited accuracy, and it is not clear how to translate the scores that they output into a likelihood of pathogenicity. The quantitative framework that we introduced here of estimating PPVs for different classes of mutations by comparing the number of observed mutations to the number expected under a well-calibrated null model of germline mutation has much more direct relevance to clinical interpretation, although the interpretation of specific DNMs still proves problematic, particularly for DNMs in sites of intermediate PPV. We propose that the scores generated by such splicing prediction tools could be calibrated by performing analogous analyses of mutation enrichment to estimate PPVs for different bins of scores. As the size of trio-based cohorts increases, the accuracy of calibration will improve.

In summary, our results demonstrate a significant contribution of non-canonical splicing mutations to the genetic landscape of DDs, a finding which is highly likely to be recapitulated across other monogenic disorders and contexts. We demonstrate the importance of non-canonical positions (particularly the don+5 site and pyrimidine-removing mutations in the PolyPy region). These inferences are supported by both population genetic investigations of purifying selection, as well as a disease based approach, considering the burden of DNMs in ˜8,000 children with severe DDs. Mutations at some non-canonical splicing positions convey a risk of disease similar to that of protein truncating and missense mutations, but are under-represented in existing databases of disease-causing variants.

## Materials and methods

### Cohort and sequencing

For full description of cohort and analytical methodology, see previous DDD publications (Deciphering Developmental Disorders 2015; Deciphering Developmental Disorders 2017). Briefly, 7,833 patients with severe, undiagnosed developmental disorders were recruited to the DDD study from 24 clinical genetics centres from across the UK and Ireland. Whole exome sequencing was conducted on the proband and both parents, with exome capture using SureSelect RNA baits (Agilent Human All-Exon V3 Plus with custom ELID C0338371 and Agilent Human All-Exon V5 Plus with custom ELID C0338371) and sequencing using 75 base paired-end reads using Illumina’s HiSeq. Mapping was conducted to GRCH37 using the Burrows-Wheeler Aligner (BWA, v0.59(Li and Durbin 2009)) and variant identification was conducted using the Genome Analysis Toolkit (GATK, v3.5.0(McKenna et al. 2010)). Variant annotation was conducted with Ensembl’s VEP (http://www.ensembl.org/info/docs/tools/vep/index.html), using Ensembl gene build 76(McLaren et al. 2016). DNMs were identified using DeNovoGear (v0.54)(Ramu et al. 2013), and filtered using an in house pipeline - denovoFilter - developed by Jeremy F. McRae(Deciphering Developmental Disorders 2017) (https://github.com/jeremymcrae/denovoFilter). Exome sequencing and phenotype data are accessible via the European Genome-phenome Archive (EGA) under accession number EGAS00001000775 (https://www.ebi.ac.uk/ega/studies/EGAS00001000775).

### Defining exons of interest

We took exons from gencode v19 (http://www.gencodegenes.org/releases/19.html) which met the following criteria: annotation_type = “exon”, gene_type = “protein_coding”, gene_status = “KNOWN”, transcript_type = “protein_coding”, transcript_status = “KNOWN”, annotation != “level 3” (automated annotation), and tag = “CCDS”, “appris_principal”, “appris_candidate_longest”, “appris_candidate”, or “exp_conf” (n = 255,812 exons)(Harrow et al. 2012). We removed a small subset of exons which no longer met these criteria in the more recent, GRCH38 based gencode v22 release (leaving 253,275 exons). We removed any exons where the median coverage at the canonical acceptor or donor positions was <15X in two sets of DDD data which used different exon capture methods (Agilent Human All-Exon V3 Plus with custom ELID C0338371 and Agilent Human All-Exon V5 Plus with custom ELID C0338371). 148,244 exons passed these criteria.

We annotated individual genomic positions relative to the acceptor and donor sites, removing any exons <14bp, and any positions which had multiple potential annotations. At the acceptor end, we considered 25bp of intronic sequence (acc-25 to acc-1) and 11bp exonic sequence (acc to acc+10). At the donor end, we considered 10bp of intronic sequence (don+1 to don+10) and 11bp exonic sequence (don to don-10). This yielded ˜6.9 million near-splice positions of interest.

The reference nucleotide composition at each position of the splicing region of interest was calculated using all sites and a weighted position weight matrix graph was generated using the seqLogo package via Bioconductor(Huber et al. 2015) (https://bioconductor.org/packages/release/bioc/html/seqLogo.html) in R (version R v3.1.3).

We define the PolyPy region as acc-3, and acc-5 to acc-17, based on pyrimidine content > 70% in our exons of interest. We assess changes from a pyrimidine to a purine (PyPu) adjusting for the strand the exon is on.

### Mutability adjusted proportion of singletons (MAPS)

In 13,750 unaffected parents enrolled as part of the DDD study, as well as >60,000 aggregated exomes from ExAC v0.3.1 (http://exac.broadinstitute.org/), we calculated the MAPS metric(Lek et al. 2016) using code developed in house by Patrick J. Short (https://github.com/pjshort/dddMAPS). The MAPS metric is based on the principle that negative selection acts to keep deleterious variation rare at a population level, but more mutable sequence contexts can contain variants that appear more common because of recent recurrent mutational events, so the metric corrects frequencies based on local sequence context using synonymous mutations. Only relevant ExAC sites with “PASS” in the VCF “FILTER” column were counted, and ExAC and DDD variants were filtered for FisherStrand (FS) <10. MAPS was calculated for all SNVs overlapping out splice positions of interest (201,587 near splice variants for DDD, and 678,241 for ExAC), the last base of the exon split by reference nucleotide (2109 variants for DDD, 6325 for ExAC), and the PolyPy, split by PyPu (15,847 variants for DDD, 58,762 for ExAC) vs all other PolyPy changes (52,300 variants for DDD, 175,287 for ExAC), as well as VEP(McLaren et al. 2016) ascertained synonymous (580,066 variants for DDD, 1,513,758 for ExAC), missense (1,125,167 variants for DDD, 2,786,533 for ExAC) and stop-gained (25,863 variants for DDD, 78,496 for ExAC) sites across autosomal regions. To establish whether the MAPS metric was significantly different between PolyPy PyPu vs all other PolyPy changes, a bootstrap resampling method was run with 1000 iterations.

### Parental variants in high pLI genes

We annotated all variant sites used in the MAPS calculations above in the 13,750 DDD parents with the gene in which the variant fell, and the pLI score of that gene, and calculated the proportion of these sites which fell within genes with high pLI scores(Lek et al. 2016) (> 0.9).

### *De novo* mutations

DNMs were identified using DeNovoGear(Ramu et al. 2013) as described in McRae *et al*, 2017(Deciphering Developmental Disorders 2017), and a stringent confidence threshold (posterior probability > 0.8) was applied. We used triplet-based mutation rates(Samocha et al. 2014) for each potential single nucleotide change across our splicing regions of interest to calculate the expected number of DNMs across autosomes in the 7,833 probands. Expected values were adjusted for depth of sequencing coverage < 50 to account for poorer ascertainment of variants in low coverage regions (exon depth <1, exp*0.119; exon depth >1 and <50, exp*(0.119+0.204*log(depth))). The values used for this correction are based on the relationship between observed and expected synonymous DNMs at different levels of coverage. We stratified this analysis into known dominant, known recessive and non-DD associated genes using the DDG2P gene list (http://www.ebi.ac.uk/gene2phenotype), downloaded in January 2016. Genes with recessive and dominant modes of inheritance were restricted to the recessive list (see Supplemental Table S4). Observed and expected numbers of DNMs were also calculated in subsets of the DDD probands with confident diagnoses (individuals with a reported variant classed as pathogenic or likely pathogenic by the referring clinician) and those lacking a potential diagnosis (diagnosed n=1417, undiagnosed n=3364, with the remainder of the cohort having possible or uncertain diagnostic states, as of January 2018). We used the Poisson test (using R’s poisson.test, with two sided alternative hypothesis) to examine differences in the observed and expected values, and a 5% FDR correction to control for multiple testing using the p.adjust R package (method=fdr) across all tests (R v3.1.3).

PPVs were calculated ((observed - expected) / observed) for CSS positions, combined and individually, don+5 sites, PolyPy PyPu, PolyPy other, other near splice exonic and intronic variants, as well as VEP defined missense and stop gained mutations.

We divided our exons into sextiles based on the pLI (Lek et al. 2016) of the gene to which they belong, and calculated the observed and expected number of DNMs in each sextile for don+5, PolyPy PyPu and synonymous variants (as above) to see if the enrichment of don+5 and PolyPy PyPu changes was concentrated in genes more likely to be intolerant of loss of function (LoF) mutations.

### Potential diagnostic variants

DNMs overlapping with our near-splice positions of interest within dominant DDG2P genes were identified in DDD probands lacking a potential explanatory variant (Dec 2016, n = 5907). The Human Phenotype Ontology (HPO, http://compbio.charite.de/hpoweb/) encoded(Kohler et al. 2017) phenotypes of the probands were assessed by consultant clinical geneticist Helen V. Firth, along with the patient’s recruiting clinician, and compared to the known clinical presentation of individuals with LoF mutations within those genes, classifying each variant as likely diagnostic, unlikely diagnostic, or unsure, depending on the strength of similarity between the proband and the disorder, and the specificity of the phenotype. The relationship between our PPVs and the proportion of clinical diagnoses in each class of near splice mutation was assessed using Pearson’s product-moment correlation using the cor.test function in R (version 3.4.4).

The proportion of CSS to non-CSS splicing diagnoses was calculated, along with 95% CIs, based on 18 non-CSS diagnoses and 48 CSS diagnoses in the same regions using the prop.test package in R (version 3.4.4).

### Validation of putative splicing variants

Eight variants were selected for validation via a minigene vector system. These comprised six likely diagnostic variants from the PolyPy, a PolyPy variant of uncertain clinical significance, and a likely diagnostic don+5 variant. Additionally, two untransmitted variants identified in unaffected parents within the same PolyPys as test variants were selected as negative controls. Details of all variants selected for validation are shown in Supplemental Table S5.

### Cloning splicing vectors

The minigene splice assay vector was adapted from that used in Singh *et al*.(Singh et al. 2016), by replacing intron 1 with the first intron from the rat insulin 2 gene (Ins-2)(Rnor_6.0 Chr1:215857148-215857695). To generate individual assay vectors, either the 5’ most 231bp (for the don+5 variant) or the 3’ most 274bp (for PolyPy variants) of this vector was replaced with the appropriate endogenous intronic sequence encompassing the DNM of interest (Figure S4a and S4b), as described below. Between 114 and 202bp flanking endogenous intronic sequence was included, along with 6bp local exonic sequence from the gene of interest.

First, proband genotypes (Supplemental Table S5) were verified by capillary sequencing of genomic PCR products (Supplemental Table S6). Genomic regions containing the reference and alternate sequences were then either amplified by nested PCR, generated by site directed mutagenesis, or generated using gene synthesis (IDT). These fragments were sub-cloned, by Gibson Assembly (NEB), into our minigene vector (Supplemental Table S7, Supplemental Table S8). The regions assayed in our vectors are detailed by genomic coordinates in Supplemental Table S5.

### *In vitro* splicing assay

HeLa cells were seeded into 12-well plates at a density of 160,000 cells per well, grown for 24 hours and transfected with 1 microgram of plasmid vector using Lipofectamine 3000 (Invitrogen). All transfections were carried out in duplicate and cultured for 48 hours. HeLa cells were cultured in DMEM (10% FCS + 1% pen/strep) at 37°C in a humidified incubator. Total RNA was extracted using a Micro RNeasy Qiagen kit and mRNA converted into cDNA using superscript IV (Invitrogen). RT-PCR was carried out using primers designed to span from exon 1 to exon 2, exon 2 to exon 3 and exon 1 to exon 3 and amplified on a thermocycler for either 25 or 35 cycles (Supplemental Table S9). Amplicons were capillary sequenced (GATC). For amplicons showing more than one splice variant (mixed capillary traces, for *CHD7*-Alt and *MBD5*-Alt), we cloned the PCR amplicons (Zero Blunt PCR cloning kit, Invitrogen) and sequenced individual colonies by capillary sequencing to identify the splice variants present (Supplemental Table S10).

Chromatograms were generated in R from .ab1 files using the sangerseqR(Hill et al. 2014) package via Bioconductor (Huber et al. 2015) (http://bioconductor.org/packages/release/bioc/html/sangerseqR.html, R v3.1.3), and likely consequences on the protein primary structure were generated using reference and alternative RNA sequences with the ExPASy Nucleotide Sequence Translation tool(Artimo et al. 2012) (https://web.expasy.org/translate/).

### Splicing pathogenicity scores

Since our region of interest spanned >6 million individual positions, each with three potential single nucleotide changes, we were restricted in the choice of splicing pathogenicity prediction tools we could utilise, as many function primarily through a low throughput web interface model. We identified three resources recently published which provide “genome wide” splicing pathogenicity scores. Two methods, dbscSNV’s AdaBoost and RandomForest are based on ensemble learning combining predictions from multiple other splice prediction tools as well as conservation and CADD scores(Jian et al. 2014a). The targeted region at the acceptor end spans 14 bases (12 intronic, 2 exonic) and at the donor end spans 11 bases (8 intronic, 3 exonic). Spidex utilises deep learning methods trained on RNA sequencing data to estimate the consequence of variants on the “percent spliced in” of an exon, relative to the reference sequence(Xiong et al. 2015). Spidex scores positions up to 300bp from intron/exon boundaries, so provides greater coverage of our splicing region of interest. We also utilised the longer standing, and widely used MaxEntScan (MES)(Yeo and Burge 2004), for which perl scripts were available, allowing the tool to be run locally for all alternative alleles of all positions of interest. The metric used for MES was the percent difference between the scores for the reference and alternative alleles, with the greatest reduction in score classed as most pathogenic. All sites were also scored with CADD(Kircher et al. 2014).

To allow cross-tool comparison, we ordered positions by increasing pathogenicity from each metric, and split positions into 20 brackets, such that the cumulative triplet based mutation rate for all variants in each bracket was equal, and the 20^th^ bracket contained the positions with the most pathogenic scores. We calculated MAPS and the proportion of parental variants falling in high pLI genes for each bracket for all five metrics, as above, and looked at the number of DNMs in known dominant genes which fell in each bracket for the five metrics. Each of these analyses was conducted including and excluding CSS dinucleotide positions.

### Splice region variants in the ClinVar database

We extracted all ClinVar(Landrum et al. 2016) (http://www.ncbi.nlm.nih.gov/clinvar/) variants using the UCSC table browser(Karolchik et al. 2004) on 02.05.2017 and matched these against our splicing positions of interest, removing exonic sites with non-synonymous consequences. This resulted in 3603 positions with clinical significance recorded as pathogenic or likely pathogenic. We calculated the ratio of canonical to non-canonical splice positions within this data. Since each variant is present in this data only once, we used number of submissions as a proxy for allele count, and calculated the ratio of canonical to non-canonical variants adjusting for this. Differences between these observed values and our expectations, based on 27% of splice affecting mutations being in non-canonical positions, were assessed using Fisher’s exact test (R v3.1.3).

## Acknowledgments

We thank the families for their participation and patience. We are grateful to the Exome Aggregation Consortium for making their data and code available. We thank the Sanger Human Genome Informatics and DNA pipelines teams for their support in generating and processing the data. We are grateful to Adam Frankish for advice selecting an appropriate exon set, and to Sarah Hunt and Fiona Cunningham for help and advice regarding splice annotation, and the development of the VEP SpliceRegion.pm plugin. Thanks also go to Alan Donaldson, Alex Henderson, Anand Saggar, Diana Baralle, Elisabeth Rosser, Elizabeth Jones, Emma Wakeling, Fleur van Dijk, Joan Paterson, Joanna Jarvis, Kate Chandler, Katherine Lachlan, Miranda Splitt, Neeti Ghali, Rachel Harrison, Sahar Mansour, Shane Mckee, Susan Tomkins and Victoria McKay for providing phenotypic information and insight on the probands with potential splice disrupting variants. The DDD study presents independent research commissioned by the Health Innovation Challenge Fund (grant HICF110091003), a parallel funding partnership between the Wellcome Trust and the UK Department of Health, and the Wellcome Trust Sanger Institute (grant WT098051). The views expressed in this publication are those of the author(s) and not necessarily those of the Wellcome Trust or the UK Department of Health. The study has UK Research Ethics Committee approval (10/H0305/83, granted by the Cambridge South Research Ethics Committee and GEN/284/12, granted by the Republic of Ireland Research Ethics Committee).

